# Transcription factor activities enhance markers of drug response in cancer

**DOI:** 10.1101/129478

**Authors:** Luz Garcia-Alonso, Francesco Iorio, Angela Matchan, Nuno Fonseca, Patricia Jaaks, Fiamenta Falcone, Graham Bignell, Simon S. McDade, Mathew J. Garnett, Julio Saez-Rodriguez

**Affiliations:** European Molecular Biology Laboratory - European Bioinformatics Institute, Wellcome Genome Campus, CB10 1SD Cambridge, UK; OpenTargets, Wellcome Genome Campus, CB10 1SD Cambridge, UK; Wellcome Trust Sanger Institute, Wellcome Genome Campus, CB10 1SD Cambridge, UK; Centre for Cancer Research and Cell Biology; Queen’s University Belfast, Belfast, BT9 7BL, UK; Joint Research Centre for Computational Biomedicine (JRC-COMBINE); RWTH Aachen University, Faculty of Medicine, MTI2 Wendlingweg 2, 52074 Aachen, Germany

**Keywords:** Transcription Factor activity, drug response, cancer heterogeneity, genomic markers

## Abstract

Transcriptional dysregulation is a key feature of cancer. Transcription factors (TFs) are the main link between signalling pathways and the transcriptional regulatory machinery of the cell, positioning them as key oncogenic inductors and therefore potential targets of therapeutic intervention. We implemented a computational pipeline to infer TF regulatory activities from basal gene expression and applied it to publicly available and newly generated RNA-seq data from a collection of 1,010 cancer cell lines and 9,250 primary tumors. We show that the predicted TF activities recapitulate known mechanisms of transcriptional dysregulation in cancer and dissect mutant-specific effects in driver genes. Importantly, we show the potential for predicted TF activities to be used as markers of sensitivity to the inhibition of their upstream regulators. Furthermore, combining these inferred activities with existing pharmacogenomic markers significantly improves the stratification of sensitive and resistant cell lines for several compounds. Our approach provides a framework to link driver genomic alterations with transcriptional dysregulation that helps to predict drug sensitivity in cancer and to dissect its mechanistic determinants.

## Background

Transcriptional dysregulation is required for tumor initiation, progression and acquisition of drug resistance[1]. Many cancer driver genes are transcription factors (TFs). Notable examples include TP53, the most commonly mutated tumor suppressor that controls cell growth arrest and apoptosis[2], and HIF1A, a key regulator of the adaptive response to hypoxic stress and the induction of angiogenesis[3]. TFs are commonly dysregulated in cancer as a consequence of a variety of genomic alterations including mutations, amplifications, deletions or chromosomal rearrangements. However, the activity of a TF can also be dysregulated through other mechanisms such as genomic alterations of their regulatory proteins. For example, HIF1A upregulation is often induced by loss-of-function (LoF) mutations in VHL[4] whereas TP53 activity can be potently suppressed through amplification of its negative regulator MDM2[5]. Due to their role as downstream effectors of signalling pathways, the aberrant activity of any protein in a pathway may ultimately result in dysregulated activity of a TF[6], which inevitably alters the expression of many of the TF’s transcriptional targets or “regulon”. Different from driver alterations in intracellular kinase-mediated signalling cascades, where redundancy may bypass the driver or provide compensatory mechanisms, aberrant transcriptional regulators have been argued to be harder to circumvent by secondary genomic alterations[7]. Consequently, TFs have been proposed as key nodal oncogenic drivers and their activity patterns used to characterise genomic aberrations in cancer[8–10] or their influence in a patient’s prognosis[11, 12].

Projects such as the Genomics of Drug Sensitivity in Cancer (GDSC)[13, 14], Cancer Therapeutics Response Portal (CTRP)[15] and the Cancer Cell Line Encyclopedia (CCLE)[16] have generated large-scale public pharmacogenomic datasets that span multiple molecular data types in a plethora of cancer cell lines. These datasets have been used to identify individual genomic, transcriptomic and epigenomic markers of drug sensitivity/resistance[13, 14, 16], thus detecting dependencies between drug response and individual molecular features. These studies recapitulated multiple clinical pharmacogenomic interactions and revealed novel possible therapeutic markers. The emerging paradigm from the aforementioned studies is one of a complex network of genomic alterations interacting with sensitivity to a large number of anti-cancer drugs. Of special interest is the potential use of these datasets to dissect the underlying molecular mechanisms regulating drug response. In order to put these informative molecular features into their operative signalling context and to shed light on the corresponding molecular mechanisms, novel and more systemic functional approaches are needed.

Here we use prediction of TF regulatory activities in cancer as sensors of pathway dysregulation. We show that the activity of a given TF can be estimated from the mRNA levels of its direct target genes extracted from DNA-binding networks, and that the activity profiles of TFs interact with genomic aberrations in upstream signalling nodes and drug response. Toward these aims, we implemented a computational framework to estimate single sample TF activity profiles across 9,250 primary tumors from the Cancer Genome Atlas (TCGA) and 1,010 cancer cell lines. For the cell lines we generated RNA-seq data for 448 cell lines, that we integrated with available RNA-seq data[16, 17]. We benchmarked different sources of TF-target interactions and computational methods to infer TF activity and assessed the prediction accuracy on independent genomics and gene-essentiality screens (Figure 1A). Then, we mined for statistical interactions between the activity patterns of the TFs and the mutational status of known cancer driver genes (Figure 1B). In order to discriminate the contributions of specific mutants, we re-annotated somatic mutations with the expected impact on the molecular properties of the coded proteins (e.g. impact on PTM regulatory sites, protein interactions, protein truncation, etc.). Finally, we investigated TF activities alone or in combination with other genomic markers as potential predictors of resistance/sensitivity to 265 of compounds (Figure 1C). Our results provide a comprehensive characterisation of TF activities in primary tumors and cell lines, show how TF activities can refine well characterised pharmacogenomic interactions and propose new testable mechanistic hypotheses on how gene aberrations influence drug response. To the extent of our knowledge, the presented study represents the first systematic evaluation of the role of TFs as markers of drug sensitivity in cancer.

**Figure 1.**
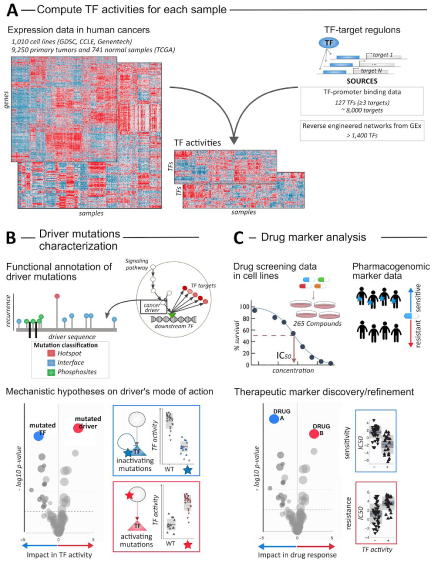
Analysis overview. A) Estimation of transcription factor (TF) activities in individual cancer samples. B) Functional evaluation of cancer mutations effect on TF activities. C) TF-based modelling of pharmacologic sensitivity, either individually or in combination with pharmacogenomic markers.

## Results

### Assembling transcriptional profiles and regulatory networks

Our initial step was to assemble a collection of basal transcriptional profiles of immortalised human cancer cell lines and primary tumors. For cancer cell lines, we extracted gene expression levels from newly derived RNA-seq data from 448 cell lines in the GDSC[13]. GDSC data was complemented with RNA-seq profiles for 934 cell lines from the CCLE[16] and for 622 cell lines from a study published by Klijn et al[17]. This collection comprises a total of 1,010 unique cancer cell line models described in COSMIC[18] (Table S1), representing, to the best of our knowledge, the largest collection of RNA-seq-derived gene expression data for cancer cell lines to date. In order to minimise technical biases introduced by different RNA-seq procedures, we processed raw reads from the three datasets using a common pipeline to derive gene-level raw counts. For primary tumors, we downloaded RNA-seq gene-level raw reads data encompassing 9,250 TCGA primary tumor samples and 741 normal samples derived by Rahman et al[19]. Raw counts from both cell lines and patient samples were further processed using a common procedure as described in the Methods section, to enable subsequent integrated analysis.

Next, we defined the set of genes whose transcription is regulated by a given TF (hereafter TF regulon). We inferred a regulatory TF-target network, through aggregating DNA-binding data from 13 publicly available resources covering TF binding site (TFBS) predictions, Chromatin immunoprecipitation (ChIP) coupled with high-throughput techniques (ChIP-X) experiments, text mining and manually curated regulatory events (Figure S1). A protein was considered a TF according to the census established by Vaquerizas et al[20]. Only TFs with at least 3 targets defined in at least two of the mentioned resources were considered. The final network consisted of 127 TFs regulating 7,978 target genes, with 106 targets per TF on average (avg) (hereafter Consensus TF Regulons, CTFR; Figure S1B; Table S2). Overall, pairwise overlap between regulons was low (avg Jaccard Similarity Coefficient = 0.0044, Figure S1C), indicating negligible levels of redundancy between most of the CTFRs.

We then used the transcriptomic data and the CTFR to derive the level of basal regulatory activity of each TF in each cancer patient and cell line based on the expression levels of their targets using the aREA algorithm in *VIPER* R package[8] (Table S3A-B). We evaluated the quality of our estimated TF activities using independent essentiality screen data from the Achilles portal (v2.4.3)[21] and Copy Number Alterations (CNA) and Whole Exome Sequencing (WES) data from the GDSC project data portal[14] (see Supplemental Results). Moreover, we studied the effect of the inclusion of CpG island methylation data to derive sample-specific CTFRs, which rendered similar results (Supplemental Results, Figures S2-S3). Finally, we compared the TF activities obtained using other regulons definition alternative to the CTFRs. Substitution of CTFRs by reverse-engineered networks from transcriptomic interactions[8] yielded slightly worse performances (Supplemental Results, Figure S2-7), hence, we decided to use CTFR to define TF-targets in the downstream analysis.

### Overview of TF activities across primary tumors, normal samples and cell lines

To obtain a global view of TFs operating in common human tumors, we studied how TF activities distribute across cancer samples. First, differential activity analysis of normal versus tumor samples over 14 tumor types revealed groups of TFs that were consistently activated or repressed across cancers. Globally, we observed a decrease in activity for the majority of TFs, whereas a small subset undergoes a strong and recurrent increase in activity across tumor types. As expected, among the upregulated TFs were the oncogenes MYC, MYCN and MAX and other genes with known oncogenic properties such as the E2F family members, FOS and FOXM1, important regulators of cell cycle gene expression; ELK1 and ETS1, involved in tumor invasion and angiogenesis[22, 23]; and HSF1, recently suggested to promote tumorigenesis[24] (Figure 2A).

**Figure 2.**
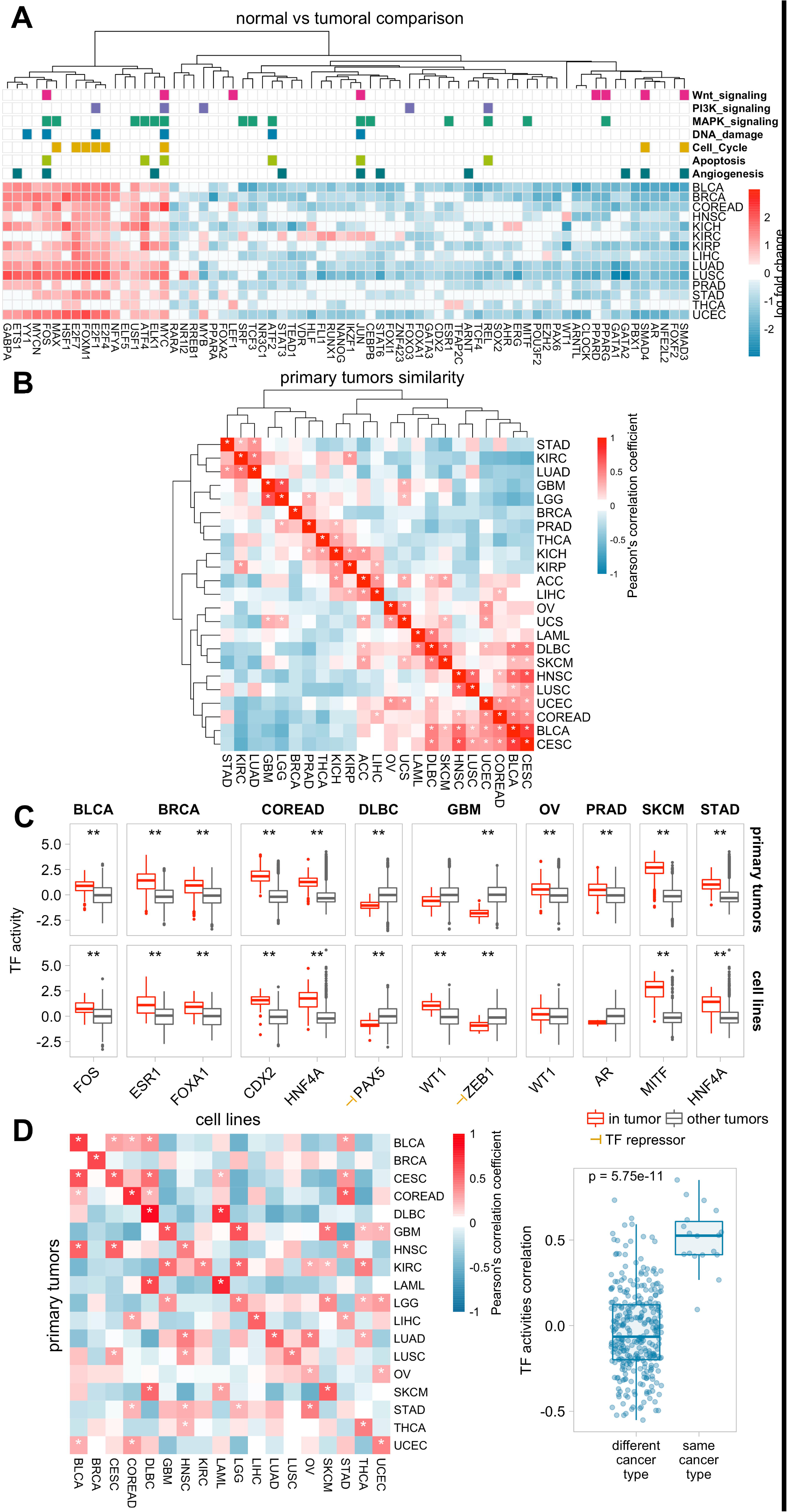
TF activities overview across primary tumors and cancer cell lines. A) Heatmap of the differential TF activity (log fold change) between normal and tumoral samples across 14 tumor types. Red and blue indicate up- and downregulation in tumors, respectively. Only TFs with a adj. p-value < 0.05 in more than half of the analysed tumors are plotted. TF-pathway associations at the top are extracted from PathwayCommons. B) Tumor type similarity. Hierarchical clustering of pearson correlation coefficients obtained from tumor type-level TF activities for 23 primary tumors. C) Activity distributions for tissue-specific TFs. Each point represents the activity of a given TF in a given sample in both primary tumors and cancer cell lines. D) Comparison of TF activities between primary tumors and cell lines for 18 common tumor types. Each value in the heatmap represents the pearson correlation coefficients between tumor type-level TF activities. Asterisks indicate significant correlations (p < 0.05).

Next, we compared the TF activity profiles between cancer types. To measure the involvement of different TFs in each cancer type, we summarised single sample-level activities into cancer type-level enrichment scores (Figure S8A-B, Table S3C-D) in both primary tumors and cell lines. Hierarchical clustering based on euclidean distance highlights similar TF activity profiles for tumors from the same tissue of origin, such as the diffuse gliomas GBM and LGG; hematopoietic and lymphoid DLBC and LAML; or squamous-like tumors such as BLCA, CESC, HNSC and LUSC (Figure 2B). These clusters are still observed in the cell line models (Figure S9). Interestingly, the TF activity profile of DLBC resembled those of cancers from the digestive system (BLCA, CESC, COREAD) and skin (SKCM) in primary tumors but not in the cell lines (Figure 2B). We hypothesised that similarities between these solid tumors and DBLC reflect the tumor immune infiltrating cells present in patient samples but not in cancer cell lines. In fact, previous studies already demonstrated different compositions of immune cells signatures across tumor types[25, 26] and suggested that subtypes of DLBC display gene expression signatures with properties similar to tumor-infiltrating lymphocytes and stromal cells[27]. However, further studies would be needed to evaluate whether these TF signatures truly reflect tumor infiltrating cell contamination.

Closer examination of well-established tissue-specific TFs (retrieved from the Human Cancer Protein Atlas[28] v15) showed that our approach captures 11 out of 12 TFs operating preferentially in specific tissues in primary tumors (Figure 2C): FOS in BLCA; ESR1 and FOXA1 in BRCA; CDX2 and HNF4A in COAD/READ; PAX5 in DLBC; WT1 in OV; AR in PRAD; MITF in SKCM and HNF4A in STAD. Note that for ZEB1, a transcriptional repressor involved in the induction of epithelial-mesenchymal transition (EMT)[29], higher protein activities correspond to a downregulation of its targets. Importantly, these tendencies are maintained in the cancer cell lines with the exception of AR, for which we observe a drop in activity in PRAD. This is in agreement with previous observations that most of the used prostate cell lines are derived from metastases and are not representative of primary PRAD[30]. These results show that our approach captures the expected activity patterns of known cancer-specific transcription factors.

A recurrent question when working with cell lines as disease models is to quantify the extent to which they mirror the molecular traits observed in primary tumors. Correlation analysis revealed an overall significant agreement (FDR < 5%) in the TF profiles between cell lines and primary samples of the same tumor type (average pairwise Pearson correlation of 0.51 and -0.04 within and between different tumor types respectively, Figure 2D), with the exception of STAD (p = 0.29, R = 0.093).

### TF activities dissect mutant-specific aberrations in cancer drivers

Previous experimental studies demonstrated that different mutations in a given protein can cause a continuum of effects, ranging from neutrality to a significant functional impact[31, 32]. We thus set-out to characterise the effect of different mutations found in well established cancer driver TFs.

As a proof of concept, we focused on TP53 due to the high frequency of mutations and their heterogeneous spectrum in cancer. We curated TP53 mutations at different levels according to: 1) specific mutation, 2) mutation hotspots 3) protein consequence 4) zygosity (only in cell lines), 5) affected domain, PTM or structural property and 6) previously proposed mutation stratifiers[33, 34] (Table S4A). Subsequently, for each of the defined groups, we compared predicted TP53 activity between mutated TP53 and wild type samples. To avoid confounding effects due to the use of samples from different tumor types, we regressed out the tissue of origin of each sample from the TF activity profiles through linear modelling. Our results indicate that all TP53 mutation significantly affecting transcriptional activity have a negative impact compared to wild type samples (Figure 3A-B; Table S4B). Overall, nonsense mutations showed a stronger impact than missense mutations as well as homozygous mutations and depletions have a stronger effect size than heterozygous mutations (Figure 3C). The mutations impacting most strongly the transcriptional activity of TP53 are the introduction of stop gain codon in position R213 and missense in positions A159 and V173. Perhaps most strikingly, comparing the three most frequent mutational hotspots R175, R248 and R273, reveals R248 and R273 mutations are amongst the most functionally disruptive, while substitutions at R175 are predicted to have lower impact in both primary tumors and cell lines. Importantly, these changes in activity are conserved between primary tumors and cell lines (R^2^ = 0.522 p = 5.8×10^−8^, Figure 3D). In order to assess if our predictions match experimental observations, we retrieved promoter-specific transcriptional measurements upon TP53 mutagenesis in yeast-based functional assays from the IARC-TP53 Mutation Database[32, 35]. Considering that variants observed in cancer samples are likely to be biased toward LoF mutations rather than neutral, our predictions on the cell lines still were in good agreement with the experimental measurements (p = 0.00749) and a similar trend, although not significant, was observed in primary tumors too (p = 0.0836, Figure 3E).

**Figure 3.**
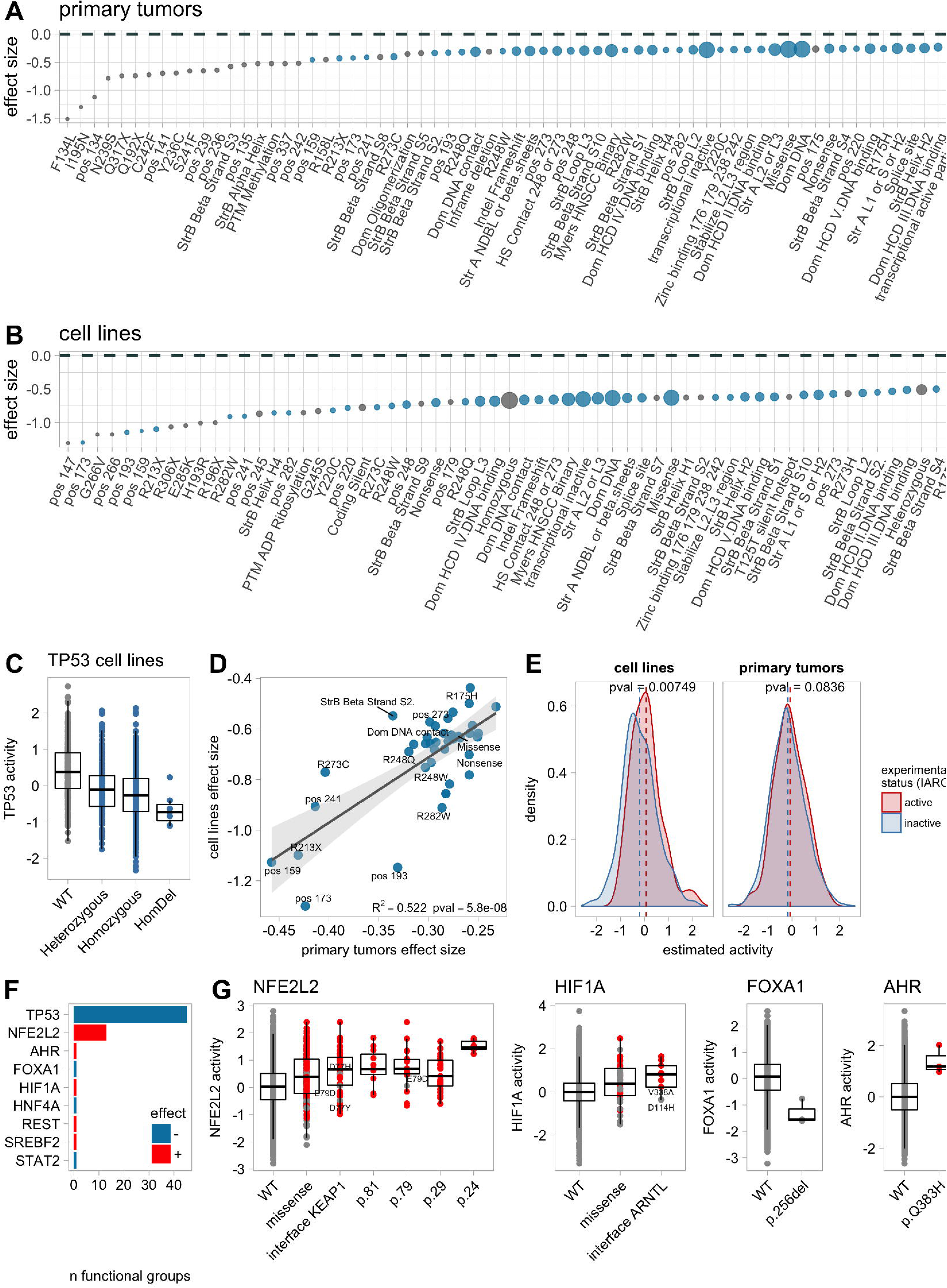
Functional characterisation of mutant TF on transcriptional activities. A and B) Effect on TF activity of different TP53 variants in cell lines and primary tumors, respectively. Y-axis indicates the effect size obtained from comparing TP53 activity between mutant and wild type samples. Negative values indicate lower TF activities in mutant samples. C) Boxplot comparing TF activities between different TP53 variants. D) Comparison of the effect size of each TP53 mutation group between primary tumors and cell lines. E) Comparison of the predicted TF activities between transcriptionally active and inactive TP53 mutants, extracted from the IARC TP53 database[35]. F) Systematic characterisation of mutant TFs in cancer samples. Each bar represents the number of mutant classes significantly affecting the activity of the mutated TF. Red and blue indicate positive and negative effects, respectively. G) Boxplots comparing TF activities across different variants of NFE2L2, HIF1A, FOXA1 and AHR, respectively.

Motivated by these results, we set out to systematically investigate the effect of the whole spectrum of mutations affecting TFs. To distinguish mutant-specific effects we studied each individual mutation and protein residue separately. Importantly, to allow us to consider non-recurrent yet potentially functional driver mutations, we also grouped mutations that, although introducing different changes in different residues, could potentially affect protein function in a similar way. Specifically, we utilised existing prediction methods and experimental reports from protein databases to group mutations according to the affected structural regions, protein interactions, post-translational modification sites, and cancer mutational hotspots (detailed description Supplemental Methods section). After aggregating those mutation classes covering the same samples (to avoid redundant groups), we recovered a total of 1250 mutation groups from 122 TFs in primary tumors.

We next evaluateed the impact of each group of mutations from our classification in the activity of the carrying TF through linear regression, pooling together samples from different tissue types. This identified a total of 9 TFs that, when mutated, exhibit a significant change in their activity profile (FDR<5%; Figure 3F, Table S4C). In general, we found that mutations in TFs with known oncogenic roles, such as NFE2L2, HIF1A and AHR were associated with increased regulatory activity, pointing to gain of function mutations. In contrast and as expected, mutations in the proposed tumor suppressors STAT2 and FOXA1 are associated with decreased regulatory activity. Also, truncating mutations in the transcriptional repressor REST resulted in increased regulon expression (Figure 3G). Analysing cell lines showed similar trends for the NFE2L2 missense mutation in D29 (p=0.009, FDR=0.0792) and REST truncating mutations (p=0.00394, FDR=0.0422).

A more detailed examination of the specific mutations responsible for such associations revealed differences in changes of protein activity associated with groups of mutations. For example, missense mutations affecting the W24/D29 residues at the surface or at the KEAP1-interface (positions 77, 79, 80, 81, 82) of NFE2L2 are associated with an increase in NFE2L2 activity, with NFE2L2^W24R/C^ mutations being associated with the strongest increase in activity (Figure 3B). NFE2L2, also known as Nrf2, is a cytoprotective factor involved in response to redox stress and genotoxic agents whose function is negatively regulated by KEAP1[36]. In cancer, NFE2L2 has been found to be recurrently mutated around the positions 24–34 and 75–82, which code the KEAP1-binding motifs DLG and ETGE respectively. Numerous studies have concluded that substitutions in DLG and ETGE motifs are positively selected as a mean to abolish KEAP1-mediated degradation of NFE2L2[37, 38].

Other examples of potential TF-activating alterations are found in HIF1A, a transcriptional regulator of the adaptive response to hypoxia. Our results suggest that samples carrying mutations at the interaction interface with its dimerisation partner ARNT-like protein (residues 42, 66, 78, 114, 245, 328, 338, 341 and 344) display increased HIF1A transcriptional activity (Figure 3G). ARNT is, together with HIF1A, another regulator of the adaptive response to hypoxia, where heterodimerisation of ARNT and HIF1A regulates HIF1A DNA binding and transactivation under hypoxic conditions[39]. To the extent of our knowledge, activating mutations in HIF1A have not been described yet.

### Driver genes regulate different TF programs

With the aim of studying how mutations in other cancer driver genes could impact the activity of TFs, we extended our analysis to cancer driver genes proposed by Vogelstein et al[40] and IntoGen platform[41]. Systematic comparisons of TF activities across mutant and wild type patient samples (considering each of 173 cancer driver genes, in turn) yielded a total number of 2,695 driver mutation groups/TF interactions (involving 146 cancer driver genes) associated with a change in activity of at least one TF in primary tumors (FDR < 5%, Figure 4A-D; Table S5A-B). The same analysis in the cell lines rendered a much lower number of associations, probably due to the lower number of samples, that involved interaction of 52 driver mutation groups/TF with 21 driver genes. Importantly, 10 of the significant mutation groups/TF associations are shared between primary tumors and cell lines with concordant effect (Fisher’s exact test (FET), odd ratio = 4.53 p < 2.8×10^−4^, Figure 4A) including RB1 and TP53 truncating mutations associated with increased activities in ATF1; KRAS mutations with increasing JUND activity; PIK3CA mutations with EGR1 activity; SMARCA4 mutations with a decrease in GATA2 activity; and EP300 mutations upregulating the activity of SMAD4 and PBX1. Some of these associations represent already proposed mechanisms of TF regulation. For example, JUND is a well known target of the ERK-MAPK pathway and its transactivation function increases upon ERK-MAPK activation[42–44]; loss of function analyses demonstrated SMARCA4 (also known as BRG1) knockout samples are defective in GATA2 activation[45]; and SMAD4 transcriptional activation to be EP300-dependent[46].

**Figure 4.**
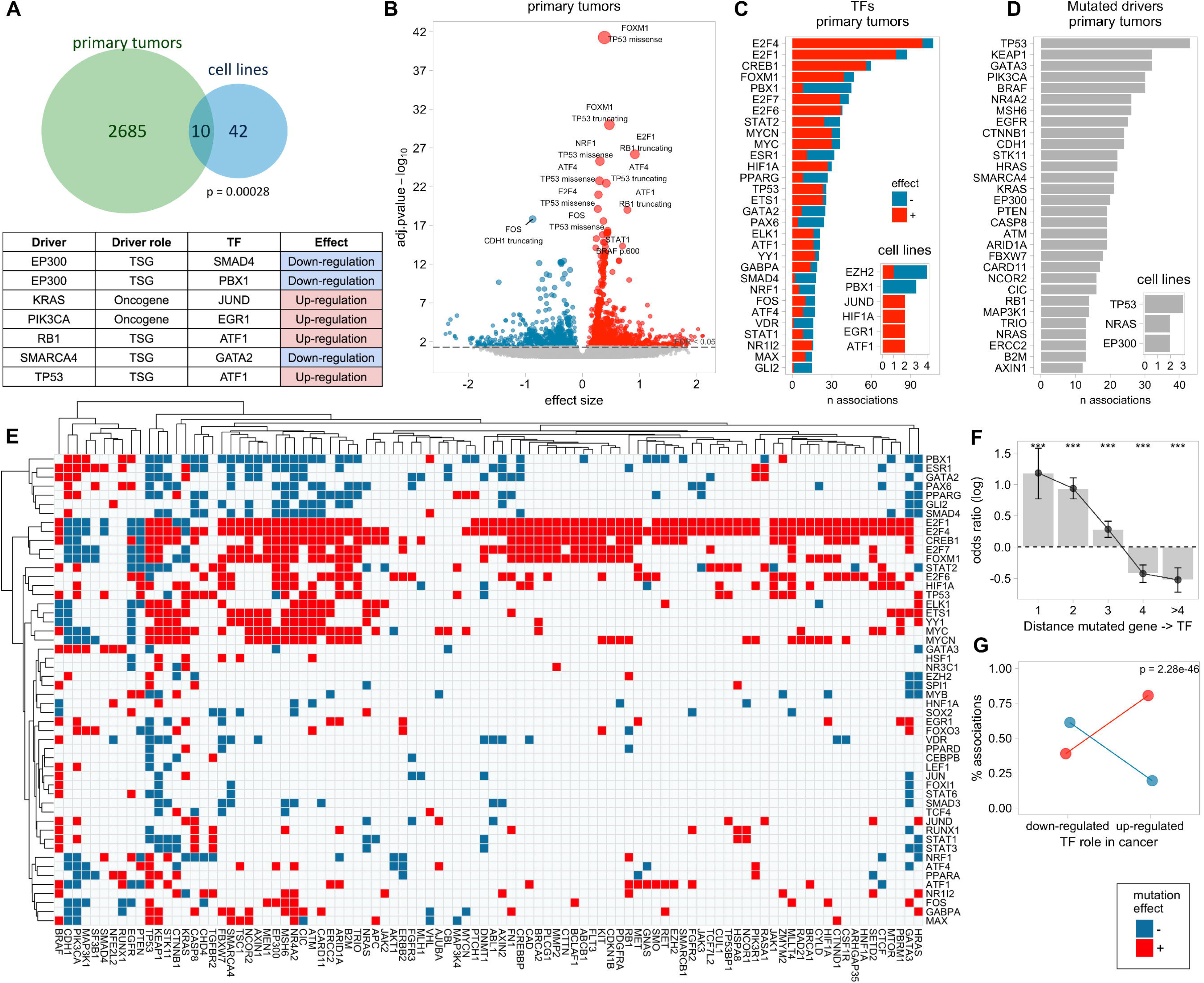
Functional characterisation of driver mutations on TF activities. A) Comparison between the TF-driver associations from primary tumors and cell lines. The Venn diagram represents total and overlapping TF-mutant significant associations. Shared driver-TF pairs are indicated in the table. B) Volcano plot with the effect size (x) and adjusted p-value of all tested pancancer associations. C) Number of significant associations per TF in primary tumors and cell lines. D) Number of significant associations per driver gene in primary tumors and cell lines. E) Heatmap of the driver-TF associations. F) Log odds ratio of finding a significant interaction by network distance (number of directed edges between the driver and the corresponding TF). G) Enrichment in positive driver-TF associations to involve oncogenic TFs and vice-versa. Colors indicate the sign of the association: red and blue correspond to significantly higher or lower TF activities in mutants compared to wild-type, respectively.

Focusing on primary tumors, the TFs predicted to be activated by a larger variety of driver genes are E2F protein members and FOXM1, key regulators of cell cycle phase transitions (Figure 4C). TP53 was the gene influencing a larger proportion of TFs (Figure 4D). Interestingly, we observed opposite patterns of effects in E2F1/4 and FOXM1 between the oncogenes CDH1 (E-cadherin), PTEN, PIK3CA, MAP3K1 and EGFR and the rest of driver genes including TP53, RB1, ATM, EP300, SMARCA4 or CREBBP, among others, suggesting that mutations in these genes tune the transcriptional machinery through different mechanisms (Figure 4E). The validity of this approach is supported by large effects size associated with RB1 suppression of E2F activity, perhaps the best described inhibitor of TF function[47], as well as the association of both ATM and TP53 in downregulating E2F and FOXM1 activity, which has recently been suggested[48–50].

In order to assess whether the detected associations represent plausible driver-TF regulatory events, we utilised the OmniPath network[51] to analyse their distance in protein-protein signalling interactions. For this purpose, we considered directed interactions and quantified the distance between every driver-TF pair in terms of shortest paths (i.e. minimum number of intermediate proteins between the driver and the TF). Enrichment analysis confirmed that significant hits tend to involve driver-TF pairs that are closer in the signalling network[51] than non-significant hits (Figure 4F). Next, we investigated whether the predicted effect of the drivers on TF activities agrees with their suggested role in cancer. We classified the TFs into 3 groups: (i) up-regulated in cancer, if the TF displays significant greater activity in tumor than in normal samples or is a known oncogene[40, 41]; (ii) down-regulated in cancer, if the TF function is repressed in tumor samples or is a tumor suppressor; and (iii) neutral, if it does not fit in any of the previous categories. Enrichment analysis revealed that positive driver-TF interactions (i.e. those representing potential TF activating events) tend to involve cancer upregulated TFs, in contrast, negative interactions are more prone to involve cancer down-regulated TFs (Figure 4G). Taken together, our results suggest that the identified associations point to likely potential mechanisms of driver-mediated transcriptional dysregulation in cancer.

### TF activity and drug sensitivity interactions in 943 cancer cell lines

We next set out to investigate the potential of the defined TF activities as markers of therapeutic response. Towards this end, we examined drug response data from the 265 compounds screened in GDSC across 943 cancer cell lines[14]. Viability reduction in response to drug treatment was expressed in terms of IC50 (drug concentration needed to achieve the half-maximal viability reduction). To identify TFs whose activity could be used as marker of drug sensitivity we made use of a linear regression approach. Pancancer and cancer-specific analyses were run in parallel, with potentially confounding factors such as the tissue of origin of the samples (in the pancancer analysis only), microsatellite instability (MI) or cell lines growth media included as covariates in our linear models[14].

The pancancer analysis identified 1,550 significant TF-drug associations (p < 0.001, FDR < 5%), with 226 out of 265 drugs (85%) and 112 out of the 127 TFs (88%) implicated in at least one interaction (Table S6A). The majority of drugs were associated with multiple TFs, which, considering the relatively low overlap in the regulons (Figure S1), may correspond to cross-correlation and functional cooperation in transcriptional regulators rather than target redundancy. We observed a large number of TF-drug associations involving relevant oncogenic TFs such as MYC, PAX5, GATA3, FOXA1, MYCN and CTCF (Figure 5A; Table S6B-C). Overall, TFs showed a tendency to interact with cytotoxic drugs (FET p < 1.77×10^−8^, odd ratio =1.6) and compounds targeting cytoskeleton, DNA replication, ERK-MAPK signalling, JNK-p38 signalling and metabolism (FET p < 0.001, Figure 5B; Table S6D).

**Figure 5.**
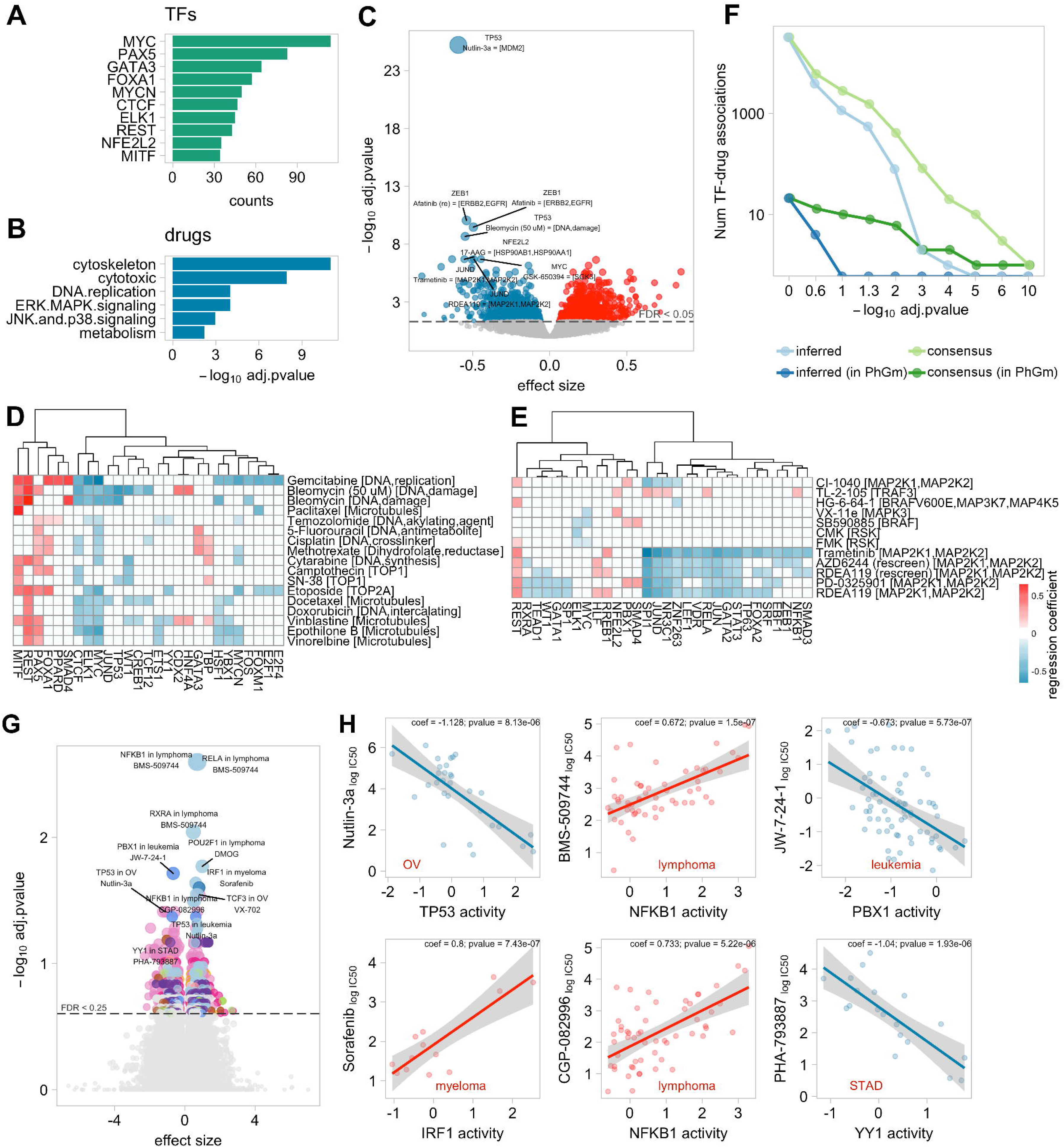
Modelling of the effect of TF activities on drug sensitivity. A) Frequency of TFs in significant pancancer TF-drug associations. B) Drug types overrepresented among significant pancancer associations. C) Volcano plot with the effect size (x) and adjusted p-value of all tested pancancer TF-drug associations. Red and blue indicate positive (resistance) and negative (sensitivity) effects, respectively. D and E) Heatmaps of significant associations with cytotoxic drugs (E) and with drugs targeting ERK-MAPK pathway. F) Number of significant pancancer associations (y) after applying different FDR cut offs (x) using consensus (green) and inferred (blue) regulons. Light lines represent all tested TF-drug associations. Dark lines represent tests involving TF-drug pairs previously defined as pharmacogenomic markers (PhGm). G) Volcano plot with the effect size (x) and adjusted p-value of all tested cancer-specific TF-drug associations. H) Examples of cancer-specific TF-drug associations. Red and blue indicate positive (resistance) and negative (sensitivity) effects, respectively.

Remarkably, the strongest detected association involved TP53 and Nutlin-3a (regression coefficient (coeff) = -0.59, p = 1.79×10^−30^, Figure 5C). Nutlin-3a is a MDM2-inhibitor that blocks MDM2-mediated TP53 degradation and enables TP53 to activate the apoptotic program. In agreement with previous studies based on mutation data, our results indicate that samples with lower TP53 activities show lower sensitivity to MDM2 inhibition[13, 14, 52]. Another strong interaction was ZEB1 upregulation, a marker of EMT, associated with resistance to EGFR inhibitor Afatinib (coeff=-0.54, p=5.32×10^−15^) and Gefitinib (coeff=-0.23, p=4.1×10^−6^). This is in agreement with a recent study in NSCLC that describes the mechanisms through which ZEB1 mediates acquired resistance to EGFR-inhibitors[53].

Reassuringly, our analyses also identified a number of groups of TFs showing simultaneous sensitivity interactions to drugs targeting common processes. For example, sensitivity to cytotoxic compounds was associated with TFs classically upregulated in actively proliferating cells (Figure 5D). In particular, sensitivity to 4 out of 5 tubulin inhibitors is associated with high MYC activity followed by ELK1, HSF1 and YBX1. Also, sensitivity to Etoposide, a topoisomerase II inhibitor, and Gemcitabine, an inhibitor of DNA synthesis, are associated with FOXM1 and E2F1/4 activities, key markers of DNA replication and cell cycle progression. Sensitivity to the two tested topoisomerase I inhibitors, in contrast, was specifically associated with CTCF and WT1 activity. Moreover, MYC and MYCN showed specific sensitivity to compounds blocking transcription elongation including CDK9[54] and RNA polymerase I inhibitors. The role of MYC in determining general sensitivity to cytotoxic agents has been controversial, with contradictory results reported in the literature. In agreement with previous studies[55–57], our results suggest that MYC-increased activity only renders cells sensitive to a subset of the tested agents, including tubulin, topoisomerases II and DNA/RNA synthesis inhibitors, but does not result in a general sensitization to cytotoxic drugs.

Focusing on targeted compounds, we noticed sensitivity associations between TFs and drugs targeting their upstream regulatory pathways. For example, sensitivity to drugs targeting the ERK-MAPK pathway (Figure 5G) was associated with increased activities in several MEK targeted TFs including SPI1, JUN, JUND and STAT3[42, 58, 59], whereas vulnerability to the two tested RSK-inhibitors correlates with ELK1 activity, another well known downstream MAPK target[58, 60]. To investigate to which extent TF activity predicts sensitivity to direct intervention of their upstream regulators, we extracted from OmniPath signalling network the proteins directly regulated by the targets of the compounds. Enrichment analysis confirmed that significant hits were more likely to involve TFs directly interacting with the drug targets (FET p = 0.0032, odd-ratio = 1.31), suggesting that predicted TF activities may be indeed indicative of upstream pathway activation and therefore useful markers of sensitivity to drugs targeting their components.

As shown, some of the investigated TFs are recurrently mutated in cancer and have already been proposed as genomic markers of sensitivity for some of the studied drugs. To validate that TF activities are, in fact, able to recapitulate drug associations with driver mutations, we compared it with the list of pharmacogenomic interactions (FDR < 25% and p < 0.001) we have previously identified for these cell lines[14]. Our approach identified 13 out of the 21 significant pharmacogenomic interactions involving a TF in our panel (FET p = 4.33×10^−4^, odd-ratio = 23.34), including TP53 mutations interacting with response to Nutlin-3a and Bleomycin; MYC with Vismodegib and PAX5 with Bleomycin.

Overall, the use of alternative reverse-engineered regulons[8] to estimate TF activities rendered fewer associations compared to CTFR on the overlapping samples: 550 (FDR < 5%, Figure S9A-B). Using an FDR threshold of 5% to define significant TF-drug interactions, these could not reproduce any of the pharmacogenomic interactions (Figure 5H), nor using a relaxed FDR threshold of 10%. Particularly, the nominal p-value for the TP53-Nutlin3a association was 0.43. Also, focusing on targeted drugs, significant hits were not enriched in TFs interacting with the corresponding drug targets (FET p = 0.99, odd-ratio=0.45).

Cancer-specific analysis revealed a lower number of associations compared to the pancancer analysis, probably due to reduced sample size (Figure 5G; Table S6E). Still, we recovered a total amount of 114 TF-drug associations (p < 0.001, FDR < 10%) including known pharmacogenomic interactions such as the TP53-Nutlin3a interaction in OV and Leukemia[52], or MYC-Temozolomide in OV[61].

Importantly, our analysis identified associations for drugs with no genomic markers reported in the cancer type under consideration (Figure 5H). Among the top hits we found that TF activity of NFKB1 (a member of NF-kappa-B complex) associated with sensitivity to ITK inhibitor BMS-509744 in Lymphoma models (coeff = 0.67, p = 7.67×10^−8^). ITK is a tyrosine kinase involved T-cell receptor (TCR) signalling pathway, whose activation triggers NF-kappa-B activity[62]. In Myeloma, resistance to Sorafenib, an inhibitor of several tyrosine protein kinases, was associated with the activity of IRF1, a proposed tumor suppressor in Acute Myeloid Leukemia[63] (coeff = 0.8, p = 5.98×10^−7^). In STAD, sensitivity to PHA-793887, a pan-CDK inhibitor, was associated with YY1, recently proposed to contribute to gastric oncogenesis[64] (coeff = -1.04, p = 1.65×10^−6^). Finally, we found sensitivity to the LCK inhibitor in Leukemia models to associate PBX1 activity (coeff = -0.66, p = 7.22×10^−7^). Aberrant upregulation of PBX1 targets has recently been reported as oncogenic factor in B-cell acute lymphoblastic leukemia (B-ALL)[65].

### TF activities enhance the predictive ability of genomic markers

We showed before that the strongest TF-drug association detected involved the well known interaction between TP53 and Nutlin-3a. According to previous studies, samples with TP53 mutations are resistant to Nutlin-3a[13, 14, 52], while our results suggest that samples with higher TP53 activities are more sensitive. We reasoned that protein activities may complement mutation-based markers and further improve the stratification of sensitive and nonsensitive cell lines. To test this hypothesis, we first confirmed that, in fact, TP53 activity was able to further identify sensitive cell lines among the wild-type samples (Figure 6A, p = 3.3×10^−16^, Likelihood Ratio test (LR)) in the pancancer context. This observation was reproduced in OV (LR p = 0.002) and a similar trend was observed in LAML (LR p = 0.1).

**Figure 6.**
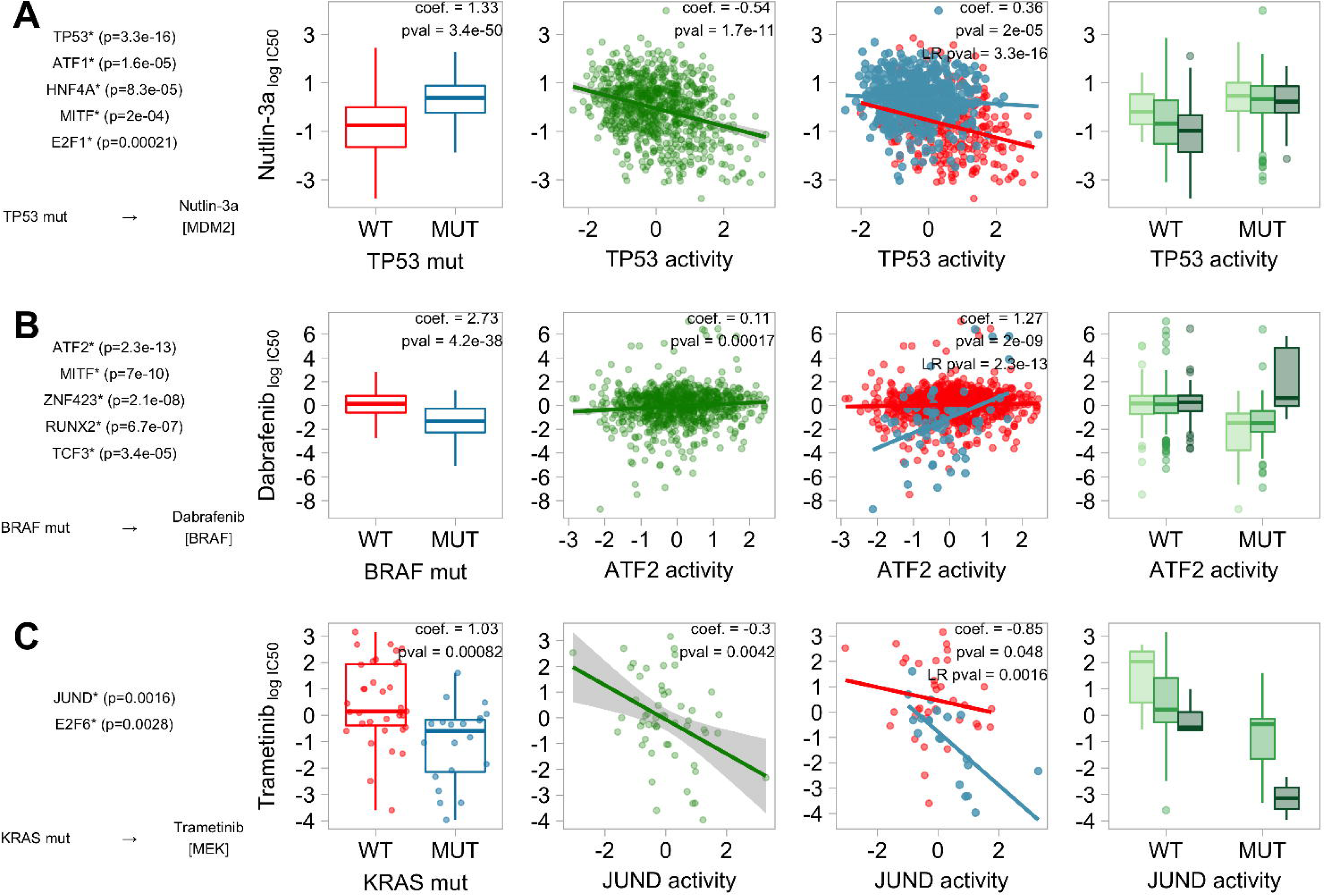
Modelling of the combined effect on drug sensitivity of known pharmacogenomic markers and TF activities. A) Top TFs whose activities enhance pancancer pharmacogenomic interaction between TP53 mutations and Nutlin-3a. B) Top TFs whose activities enhance pancancer pharmacogenomic interaction between BRAF mutations and Dabrafenib. C) Top TF whose activity enhances pharmacogenomic interaction between KRAS mutations and Trametinib in LUAD. First boxplot represents the IC50 (y) of an individual cell line in mutant (blue) and WT (red) samples. The second scatterplot represents the relationship between the IC50 (y) and the predicted TF activity (x). The third scatterplot represents the relationship between the IC50 (y) and the predicted TF activity (x) in mutant (blue) and WT (red) samples.

Motivated by this finding we ran a systematic analysis to search for TFs able to refine known pharmacogenomic interactions. Overall, we observed that 86 out of 158 (54.4%) tested strong effect pharmacogenomic interactions identified in Iorio et al., 2016[14] are improved by at least one TF (FDR < 5%, LR test; Table S7). Again, we observed groups of TFs interacting with the same drug/pharmacogenomic marker. Significant hits are strongly enriched for TFs improving pharmacogenomic interactions involving targeted compounds (FET p=1.36×10^−33^), particularly compounds targeting Receptor Tyrosine Kinases (RTKs), ERK-MAPK and PI3K pathways (FET p < 0.001).

The second strongest hit after TP53-Nutlin3a involved the interaction between BRAF mutational status and the FDA approved BRAF inhibitor Dabrafenib. Specifically, in mutant BRAF samples, resistance to Dabrafenib interacts with ATF2 and MITF regulons (Figure 6B, p = 2.34×10^−13^ and p = 6.98×10^−10^), this last one a marker of skin cells. Resistance in BRAF mutants to Dabrafenib was still observable in SKCM samples with higher expression of ATF2 targets (p = 0.0013). The importance of ATF2 in melanoma is supported by several lines of evidence; ATF2 is required for melanoma tumor development[66]; nuclear ATF2 (transcriptionally active) is associated with poor prognosis, metastasis and resistance to genotoxic stress; mitochondrial ATF2 (transcriptionally inactive) is associated with increased apoptosis[67, 68]. Moreover, PKCε, the kinase mediating ATF2 transcriptional activity, is among the top 10 kinases associated with BRAF-inhibition resistance, which supports the relationship between ATF2 and Dabrafenib resistance[69]. Finally, this observation is also supported by gene-level essentiality scores from Achilles project, where we found a pancancer tendency between the predicted activity for ATF2 and its essentiality in BRAF^V600E^ mutant cells (R = -0.55, p = 0.062, Pearson correlation; Figure S10A) but not in BRAF^wt^ (R = 0.078, p = 0.38, Pearson correlation; Figure S10B).

Interestingly, the most significant improvements in predictions were observed between drugs targeting ERK-MAPK signalling (FET p=5.36×10^−8^) and the driver genes BRAF, KRAS or HRAS. For example, in BRAF wild type samples, sensitivity to MEK inhibitors improved including JUND in the model, among others (p = 4.56×10^−11^ and p = 4.71×10^−11^, RDEA119 and Trametinib respectively). Our previous analysis already suggested JUND regulon to be predictive of MEK-inhibition sensitivity alone. Here we show how JUND also improves response prediction to MEK inhibitor AZD6244 within HRAS mutant pancancer samples (p = 1.9×10^−6^) and to Trametinib in within KRAS mutated LUAD samples (p = 1.4×10^−3^, Figure 6C). Several studies have already proposed JUND as a downstream substrate of the ERK-MAPK signalling pathway[58, 70]. Taken together, our results suggest that JUND regulon may be used as a sensor of ERK-MAPK pathway activity and vulnerability to MEK-inhibition.

Finally, other potential interactions affecting well established pharmacogenomic markers are: the interaction of JUND with sensitivity to cell cycle CDK4/CDK6 inhibitors in samples with RB1 mutations (p=1.9×10^−6^), which in turn is known to regulate cyclins[71, 72]; sensitivity to AKT inhibitor[73, 74] GSK690693 interaction with several TFs in OV PIK3CA mutated samples, where the strongest hit involves CREB1 (p=2.2×10^−6^), the key downstream effector of the PI3K/Akt/CREB signalling pathway[75]; and sensitivity interaction between ERBB2 inhibitors Lapatinib and CP724714 with activity of ELF1 in HER2+ BRCA samples (p=1.1×10^−5,^ p=2.1×10^−5^), a candidate regulator of ERBB2 expression[76].

## Discussion

TFs activities derived from gene expression data have attracted much attention in cancer research during the last few years. Recent studies have applied DNA-binding networks derived from ENCODE ChIP-Seq data to compare TF activity profiles across different cancers and evaluate their potential as prognostic markers[11, 12]. Alternative approaches have estimated protein activities using tumor-specific inferred gene networks and applied them to characterise the impact of somatic alterations[8, 9], proposing new hypotheses on how specific driver mutations may alter transcriptional regulators. Although based on different definitions of TF regulons, the common outcome is that the estimation of transcriptional activities from mRNA levels of TF targets can reveal novel mechanisms involved in tumor development. However, the potential use of TF activity as markers to guide personalised treatments, alone or in combination with established genomic markers, has not yet been explored.

Here, we applied an unsupervised analysis pipeline to derive signatures of TF activity from new and existing RNA-seq data in 1,010 cancer cell lines and 9,250 primary tumors. Our approach circumvents the need to turn to prior classification of samples into subtypes, of particular benefit when working with heterogeneous group of cancer patients, and avoids the use of a reference unperturbed control for systematic comparisons, which is not always available, specifically for cancer cell lines. These TF signatures enabled us to (i) functionally characterise different mutations impacting driver TFs; (ii) link genomic aberrations in drivers with TF dysregulation; (iii) suggest new mechanisms for response to specific compounds in cancer models and (iv) propose new markers of drug response, alone or in combination with genomic markers. To our knowledge, the results obtained provide the first systematic exploration of interactions between TF activities and drug response in cancer. Although we expect some interactions to reflect the cooperative behaviour between TFs controlling common processes rather than causal associations, we found that these recapitulated known pharmacogenomic relationships and were enriched for TF-drug pairs where the targeted genes were close upstream in the signalling network to the associated TF. Thus, we envision that the identified associations provide reliable evidence to refine existing hypotheses or formulate new ones to understand therapeutic outcomes. Finally, our study shows that predictions on therapeutic response can be improved if, in addition to the mutational status of a genomic marker, the regulatory activity of the involved protein is also considered. This can be achieved directly, when the gene marker codes for a TF as exemplified by TP53 -Nutlin3a response, or indirectly, when the coded protein regulates a TF as the case of JUND in MEK inhibitors.

The critical factor in the quantification of TF activities is the definition of the targets putatively regulated. Here, we chose to use a curated compendium of regulatory networks derived from different TF-DNA binding evidences, such as *in vivo* ChIP-X experiments, *in silico* TFBS predictions and literature-based collections of regulatory interactions, that we called consensus TF regulons (CTFRs). The major limitations of our approach are (i) the incomplete knowledge of the targets belonging to each TF regulon, (ii) the assumption that a TF either induces or represses its targets (but TFs may act as both activators or repressors of gene expression) and (iii) the contextual-dependencies of the sources of TF regulons[77]. In the light of these considerations, approaches inferring condition-specific regulatory networks from transcriptional interactions have become very popular in the last decade[78]. The underlying principle of most methods is that TF circuits can be inferred through correlating mRNA levels of the TFs with all other genes[79, 80]. However, this assumes that the mRNA expression level of a gene is a good indicator of the activity of the corresponding protein, which applies in some cases but may fail for TFs whose activity depends on post-transcriptional, post-translational regulation, modifications (phosphorylation, acetylation etc.) or indeed their stoichiometric assembly in a range heteromeric complexes[81]. Moreover, an additional important phenomenon disrupting this assumption, also supported by the findings presented herein, is the pervasiveness of cancer related mutations in TFs that change the protein function. Pertinent examples are LoF TP53 missense mutants, which while abundantly present at mRNA and protein level, are predominantly unable to directly regulate the expression of its canonical targets. Furthermore, these methods are susceptible to be confounded by indirect associations or co-expression of other TFs[82]. Finally, the inference of such condition-specific networks require a prior classification of samples, which may not be trivial for heterogeneous cancer cell line panels.

Nonetheless, our TF predictions based on CTFRs agree with independent essentiality screenings and genomic data and mimic changes in transactivation potential observed in mutagenesis studies. Importantly, CTFRs are able to reproduce known pharmacogenomic interactions while inferred regulons fail to do so. However, it is worth mentioning that our strategy to retrieve CTFRs may favour well studied TF, whose targets have been thoroughly characterised, thus resulting in biased performances. Further refinement of the approaches to define TF regulons and approximate their activity in cancer should enable to find further pharmacogenomic interactions and thereby novel markers and therapeutic opportunities.

## Conclusion

A major challenge in cancer research is the stratification of patients for therapeutic intervention. Although, to date, the majority of the strategies focus on the identification of somatic genomic aberrations as predictive response factors for anti-cancer therapies, there is still a plethora of cancer subtype therapies for which known driver aberrations alone have failed to show any predictive ability. Here we investigated the basal activity of TFs in 1,010 cancer cell lines and 9,250 primary tumors derived as a proxy of the expression of their high confidence target genes. To the best of our knowledge, this study represents the largest functional evaluation of basal TF activities integrating cancer genomics and drug response data to date. Our results demonstrate that TF activity profiles derived from curated TF-DNA binding data can be used to characterise genomic alterations and drug response in cancer patients, proposing these as promising complementary markers of therapeutic response. The proposed approach may have strong implications in the refinement of personalised treatment methodologies. We envision that with the increase in the coverage and quality of the CTFRs, the proposed strategy will become instrumental to interpret transcriptional dysregulation in cancer and elucidate its clinical implications.

## Methods

### Cell lines and primary tumors data

RNA-seq data: RNAseq data for 448 cell lines were sequenced in-house (EGAS00001000828). RNA libraries were made with the Stranded mRNA library prep kit from KAPA Biosystems according to the KAPA manual using the Agilent Bravo platform. For the rest of cancer cell lines, fastq files were downloaded from CCLE[16] (PRJNA169425) and Klijn et al[17](EGAS00001000610). The iRAP pipeline [83] was used to filter low quality reads, alignment and raw counts quantification of the three cancer cell lines RNA-seq datasets. Annotation and genome reference was based on Ensembl release 79. For TCGA samples, raw counts derived from an alternative processing pipeline were directly downloaded from the Gene Expression Omnibus at the accession GSE62944[19]. Raw counts from cell lines and patients where processed independently but using a common pipeline, to maintain consistency, as recommended by *limma* protocol prior to a *voom* transformation[84]. Briefly, 1) samples which proportion of genes with 0 counts[85] exceed 40% were discarded; 2) lowly expressed genes, defined as those with an average CPM lower than 1, were discarded; 3) data was normalised using TMM approach described in *edgeR* package; 4) a *limma-voom* transformation was applied to the data and fitted log2 counts per million with associated precision weights were extracted. Finally, for cell lines, data was batch corrected using *ComBat* from *sva* R package[86] to account for the possible bias effects introduced by different platforms in the experimental protocols used to generate the RNA-seq data (GDSC, CCLE and Klijn et al), keeping the tissue of origin as a covariate.

WES data: For cancer cell lines, we used the list of genomic variants assembled from the COSMIC database available through the GDSC1000[14, 87, 88]. For TCGA primary tumors, we downloaded WES data from the cBioportal[89]. All genomic coordinates of variants refer to human genome assembly GRCh37. To maintain consistency in the annotations between both datasets, genomic coordinates of WES variants were re-annotated with ANNOVAR version 2.4[90] and mapped to ensembl gene coordinates under the genome build version hg19. The final datasets contained a total of 608608 and 982200 WES variants for cell lines and primary tumors, respectively.

CNA data: For cancer cell lines, we downloaded PICNIC[91] processed data from the GDSC1000[14, 87, 92]. For TCGA primary tumors, CNA GISTIC[93] scores were downloaded from the cBioportal[89]. A gene was considered to be homozygously depleted if the maximum copy number of any genomic segment containing coding sequence of the gene from PICNIC was 0, in the case of the cell lines, or the GISTIC score was equal to -2, for the primary tumors.

Drug response data: Effects on cell viability for 265 compounds in the cancer cell lines were downloaded from the GDSC1000 data portal[14, 87, 94]. Dose-response was defined as the natural logarithm of the half-maximal inhibitory µM concentrations (IC50).

Methylation data: For the cell lines, information on the methylation status of the promoter regions of coding genes was downloaded from the GDSC1000 portal[14, 87, 95]. Specifically, the downloaded data represents binary events referring to low and high methylation status of CpG islands derived from per gene averaged beta values in gene promoters.

Clinical data: For cell lines, annotation on cancer types (GDSC.description_1 and GDSC.description_2), TCGA identifier, microsatellite instability status, growth properties and media was downloaded from the GDSC1000 portal[14, 87]. For primary tumors, information on TCGA cancer type identifier and clinical variables was downloaded from cBioportal[89].

Gene essentiality data: We downloaded ATARiS phenotype values (version v2.4.3), reflecting the relative effects of gene suppression across 216 cell lines, from the Achilles portal[21, 96]. A gene was defined to be essential in a sample if the ATARiS value was 2 standard deviations away from the mean value of the same gene across the whole population of cell lines. Additionally, genes deviating 3 standard deviations away from the mean were defined as highly essential.

Table S1 summarises the number of samples covered by each data type and the overlap with respect to the expression data.

### TF regulons data

We built two types of TF regulatory networks. The first type, called Consensus TF Regulons (CTFR), was derived by aggregating TF-target regulatory interactions from different publicly available sources of TF-binding evidences: TFBS predictions (FANTOM[97], JASPAR[98] and TRANSFAC via MSigDB[99]), ChIP-X data (ChEA[100], HTRI[101]), literature text-mining (TRRUST[102]) and manually curated interactions (KEGG[103] and ORegAnno[104]). A TF-target interaction was included in the final regulon if it was defined in at least two of the mentioned resources. A TF regulon was only used in the analysis if it contained at least 3 targets with measured expression data. TF-target interactions in this type of regulon were unsigned and weighted equally.

The second set of TF regulons was derived from inferred gene networks[80] downloaded through *aracne.networks* R package in Bioconductor. Here, reverse engineered networks were built on a cancer-specific way from 24 TCGA tumor datasets as described in[8]. Cancer-specific TF-target relationships were weighted and signed, where weights were derived by ARACNe through a mutual information approach and signs were derived from the spearman correlation coefficient between the TF and the mRNA levels of the corresponding target gene.

In both network types, a protein was defined to be a TF if it was classified as such in the census proposed by Vaquerizas and colleagues[20] or contains the keyword “*transcription factor*” in the Uniprot database[105]. Unlikely TFs, as defined by Vaquerizas and colleagues, were discarded out from TF census. CTFR regulons are available in the Table S2.

### Scoring basal activity of TF

The input of the method is a matrix of normalised expression values for *N* samples and *M* genes. The first step consists in the gene-wise normalisation of the expression distribution by using a kernel estimation of the cumulative density function (kcdf)[106]. Therefore, these gene expression estimates are relative to the population under study. Next, under the assumption that TFs function by modulating the transcription of their target genes in a coordinated way, the level of activity of a TF in a sample was approximated as a function of the collective mRNA levels of the TF’s target genes using the aREA algorithm from the *VIPER* R package. Positive scores indicate relatively greater basal activity of the TF in the sample compared to the background population, whereas negative scores indicate lower basal activity or repression.

### Normal-tumoral comparison of TF activities

Differential TF activities between normal and tumoral TCGA samples were computed using the limma R package. Specifically, the matrix of normalised TF activities per sample was used to fit a linear model for each TF (lmFit) and the empirical Bayes (eBayes) test was used to obtain the corresponding moderated t-statistics together with the nominal and adjusted p-values[84]. An independent test was run for each cancer type.

### Primary tumor-cell line comparison of TF activities

In order to summarise single sample-level into cancer type-level activities, we used *VIPER* to compute activity enrichment scores per TFs and cancer types. Next, pearson correlation was used to compare the vector of activities from the cell lines with the vector of activities from the primary tumors. For comparison purpose, we used the corresponding TCGA label mapping in the cell lines (available at GDSC data portal).

### Statistical models of TF activity association with drug response

We used a linear model (no interaction terms) per drug to correlate response with TF activities. Here, for each drug-TF pair, a vector of IC50 values per sample was modeled as a function of the dependent variables[14] defining tissue type (only for pan-cancer analyses), microsatellite instability status, the screening medium factors and the estimated activity of the TF. These factors were shown to influence the response to several compounds and are added into the model to account the possible confounding effect. The impact of the TF on the drug response, that is the relative difference in the mean IC50 according the variation in the TF activity, was defined by the magnitude of the regression coefficient estimated by solving a multiple linear least squares regression. The type-II analysis-of-variance (ANOVA) from the *car* R package was used to calculate a F-tests and obtain the significance of the regressors. Finally, for each cancer type, p-values were adjusted for multiple testing correction using Benjamini Hochberg method.

In order to include as many cell lines as possible, tissue factors in the pan-cancer models were defined by the GDSC.description_2 due to the presence of a significant amount of cell lines without a matching TCGA type. For cancer specific analyses, TCGA labels were used for consistency with GDSC1000 study on pharmacogenomic markers[14]. Moreover, to cover a broader spectrum of tumor types in the cancer-specific study, we also run independent analyses in Ewing’s sarcoma, leukemia, lymphoma, osteosarcoma and rhabdomyosarcoma samples as defined in the GDSC.description_1.

### Statistical contribution of TF activities in improving known pharmacogenomic markers

Pharmacogenomic associations were extracted from the GDSC1000 portal[14, 87]. Only statistically significant large-effect interactions (p < 0.001, FDR < 20% and Glass Δs > 1) were considered. Genomic markers occurring in less than 3 cell lines in our final population were discarded.

To evaluate the extent to which TF activities can improve the predictive power of known genomic markers alone on drug-response, we compared linear regression models with and without the TF activity as a dependent variable as an interaction term with the genomic marker. Models were compared using a log likelihood ratio test. Resulting p-values were adjusted for multiple testing correction. A 5% FDR threshold was used to define models with TF activities to fit significantly better than the corresponding models without the TF predictor.

### Functional annotation of driver mutations and classification

We classified WES variants in driver genes by estimating the biological implications of each alteration on the protein. Consequences of short deletions, insertions, and nonsense mutations were classified as protein truncating. In contrast, consequences of missense mutations can be broader. To estimate their possible consequences we applied a series of mapping strategies and publically available tools accounting for the location of the mutation within the protein structure, regulatory sites, cancer mutational hotspots, the impact in activity, protein stability and their affinity to associate, recognise or be recognised by other molecules. See a detailed description in the Supplementary Methods section.

Vogelstein et al[40] and IntoGen[41] census were used to define genes that are cancer drivers. Moreover, we added in the analysis potentially new driver genes identified by eDriver[107] and ActivDriver[108] and the TFs under study.

### Statistical models of drivers association on TF activities

Similarly, we used ANOVA (no interaction terms) per TF to correlate protein activities with the mutational status/class of driver genes. Here, for each TF and each group of driver mutations, a vector of TF activities was modeled as a function of the dependent variables defining tissue type (only for pan-cancer analyses), microsatellite instability status, the screening medium factors and the status of the driver (wild type or mutated). Only genomic markers occurring in at least 3 cell lines in the studied population were considered. The effect of the mutations on the measured TF activities with respect to the wild type was defined by Cohen’s d effect size estimation. A type-II ANOVA from the *car* R package was used to obtain the significance of the regressors. Finally, p-values were adjusted for multiple testing correction by FDR method on a cancer type basis.

## List of abbreviations

ACC: Adrenocortical carcinoma
ALL: Acute myeloid leukemia
BLCA: Bladder carcinoma
BRCA: Breast carcinoma
CCLE: Cancer Cell Lines Encyclopedia
CESC: Cervical squamous carcinoma
ChIP: Chromatin immunoprecipitation
ChIP-X: Chromatin immunoprecipitation coupled with high-throughput technique
CNA: Copy number alteration
COREAD: (COAD/READ) Colon adenocarcinoma/Rectal adenocarcinoma
CTFR: Consensus transcription factor regulon
DLBC: Difuse B cell lymphoma
EMT: Epithelial-mesenchymal transition
FDR: False discovery rate
FET: Fisher’s exact test
GBM: Glioblastoma multiforme
GDSC: Genomics of Drug Sensitivity in Cancer
HNSC: Head and neck squamous cell carcinoma
KICH: Kidney chromophome
KIRC: Kidney renal clear cell carcinoma
KIRP: Kidney renal papillary carcinoma
LAML: Acute myeloid leukemia
LGG: Lower grade glioma
LIHC: Liver hepatocarcinoma
LUAD: Lung adenocarcinoma
LUSC: Lung squamous cell carcinoma
MB: Medulloblastoma
MM: Myeloma
NSCLC: Non-Small cell lung carcinoma
OV: Serous ovarian adenocarcinoma
PRAD: Prostate adenocarcinoma
SKCM: Skin carcinoma
STAD: Stomach adenocarcinoma
TCGA: The Cancer Genome Atlas
TF: Transcription factor
TFBS: Transcription factor binding site
THCA: Thyroid carcinoma
UCEC: Uterine corpus endometrioid carcinoma
WES: Whole exome sequencing

## Additional files

**Additional file 1:** Supplemental results, materials and Figures S1 to S10.

**Additional file 2: Table S1.** Summary of number of samples per dataset used in this study.

**Additional file 3: Table S2.** TF-target interactions in the CTFs.

**Additional file 4: Table S3.** TF activities using CTFRs. Sample-level activities in (A) primary tumors and (B) cell lines. Summaries of cancer-level activities in (C) primary tumors and (D) cell lines.

**Additional file 5: Table S4.** Functional characterisation of TF mutations. (A) Manual classification of TP53 mutations. (B-C) Full list of ANOVA pan-cancer interactions between TP53 mutation classes and TP53 activity in primary tumors and cell lines, respectively. (D-E) Full list of ANOVA pan-cancer interactions between TF mutation classes and their corresponding activity in primary tumors and cell lines, respectively.

**Additional file 6: Table S5.** Functional characterisation of driver mutations. (A) Full list of ANOVA pan-cancer interactions between mutation classes in driver genes and TF activities in primary tumors and (B) cell lines, respectively.

**Additional file 7: Table S6.** TF-drug association analysis. (A) Full list of pan-cancer interactions between TF activities and drug response. (B) Frequency of each TF in the significant associations. (C) Frequency of each drug in the significant associations. (D) Enrichment of drug classes among the significant hits. (E) Full list of cancer-specific interactions between TF activities and drug response.

**Additional file 8: Table S7.** Refinement of pharmacogenomic interactions analysis. Full list of pan-cancer interactions between TF and string effet pharmacogenomic interactions identified in Iorio et al 2016.

## Declarations

Ethics approval and consent to participate

Not applicable.

### Consent for publication

Not applicable.

### Availability of data and material

R code to compute TF activities is available in the github repository: https://github.com/saezlab/DoRothEA

The datasets generated and/or analysed during the current study are included in this published article and its supplemental information files. Processed RNA-seq data is available from the corresponding author on reasonable request.

### Authors’ contributions

Conceived and designed the experiments, L.G.-A., F.I. and J.S.-R; Methodology and analysis, L.G.-A. and F.I.; RNA-seq processing, L.G.-A., A.M., N.F. and G.B.; Data curation, L.G.-A., F.F. and S.M.; Interpretation of results, L.G.-A., F.I. and P.J.; Writing – original draft, L.G.-A, F.I. and J.S.-R.; Writing – review & editing, L.G.-A, F.I., P.J., S.M., M.J.G. and J.S.-R.; Supervision, J.S.-R.; Funding acquisition, M.J.G and J.S.-R.

## Acknowledgements

We thank the Gene Expression Atlas team for the help with the RNA-sequencing processing, specially Laura Huertas for the curation of sample annotations and Robert Petryszak and Alvis Brazma for the general support. We thank Cyril Benes, Pedro Beltrao, Ivan Costa and Ian Dunham for insightful discussions and providing valuable feedback on the manuscript; and Euan Stronach, Paul Fisher and Glyn Bradley for input in design and analysis.

